# Computational Screening of Filamin Mechanical Binding Proteins using AlphaFold2

**DOI:** 10.1101/2025.05.02.651884

**Authors:** Jennifer Johnson, Nicanor González-Morales

## Abstract

Filamins are dimeric actin binding protein that play a critical role in mechanical signaling. They contain a mechanosensory region (MSR) that naturally folds into a globular closed conformation. Under mechanical stress, the MSR unfolds into an open conformation, exposing binding sites for numerous proteins. Filamins are involved in diverse cellular functions, and their mechanical binding targets are highly context dependent. In this study, we employed AlphaFold2 modeling for screening proteins that specifically recognize the open conformation of filamins. We focused on the *Drosophila melanogaster* filamin, Cheerio, and conducted a biased screen to identify mechanical binding proteins. We selected the top 132 hits from the initial screening for further characterization. All identified binding proteins specifically recognize the open conformation of the MSR and not the closed conformation. Interestingly, the binding regions of these proteins lack obvious sequence similarity. While some false positives were identified, they could be effectively filtered out based on the secondary structure formed at the binding interface. This study provides a framework for identifying specifically filamin interactions in mechanosignaling.

## 1 Introduction

Filamins are a family of large cytoskeletal proteins that play a crucial role in the organization of the actin cytoskeleton in cells, primarily by sensing the stretching of the actin cytoskeleton [1–3]. Typical filamins are composed of an N-terminal actin-binding domain, a series of immunoglobulin (Ig) domains responsible for mediating protein-protein interactions central rod domain, and a C-terminal dimerization domain [3]. The Ig domains 14-19 in *Drosophila* and 16-21 in vertebrates constitute the mechanosensitive region (MSR), which adopts a globular organization that underlies the force-sensing role of filamins [2,4,5]. At a resting state, the MSR adopts a closed conformation that masks binding sites located in Ig domains 17,19, and 21 in vertebrates or 15,17, and 19 in *Drosophila* [4–8]. When filamin is stretched by changes in the actin cytoskeleton the MSR unfolds revealing specific protein binding sites. The last Ig domain constitutes the dimerization domain [9–11]. Signaling proteins which recognize the unfolded MSR provide feedback to the actin cytoskeleton network for it to adapt. *Drosophila* has a single typical filamin protein called Cheerio (Cher), which shares the structure and function of typical filamins [3,12].

The Ig domains of the MSR region consist of seven β-strands (A-G) that form an immunoglobulin-like β sandwich [1,6–8]. The Ig domains of the MSR have a ligand-binding site on the C to D β-strand face. When a ligand binds, an additional anti-parallel β-strand is formed next to the C β-strand [1,6,8]. The ligand residues contribute to the β-strand and form hydrogen bonds, while specificity is determined by side-chain interactions with the groove between Ig β-strands C and D. Ligand residues interact with the groove via their side chains through hydrogen bonds and hydrophobic interactions. The Ig domain-binding motif is a typical β-strand-forming sequence with alternating hydrophobic residues pointing toward the interior of the protein and the CD face appears to be a general mode of interaction for filamin ligands such as migfilin[13,14], FilGAP[15,16], CFTR[17], and integrins [8,18]. At resting state, the MSR is in a closed form, where Ig 16 blocks the C β-strand of Ig17 and Ig 18 blocks the C β-strand of Ig19 [6,8]. The closed conformation blocks the associations with other peptides, which also use the C β-strand face (Figure 1). Upon a pulling force, the β-strand from the blocking domains is released exposing a binding site in the C β-strand of Ig17 and Ig19 [5]. β-strand unmasking occurs at forces of 2-5 pN and Ig domains of filamins are stable at pulling forces of less than 35pN [19,20].

**Figure 1.**
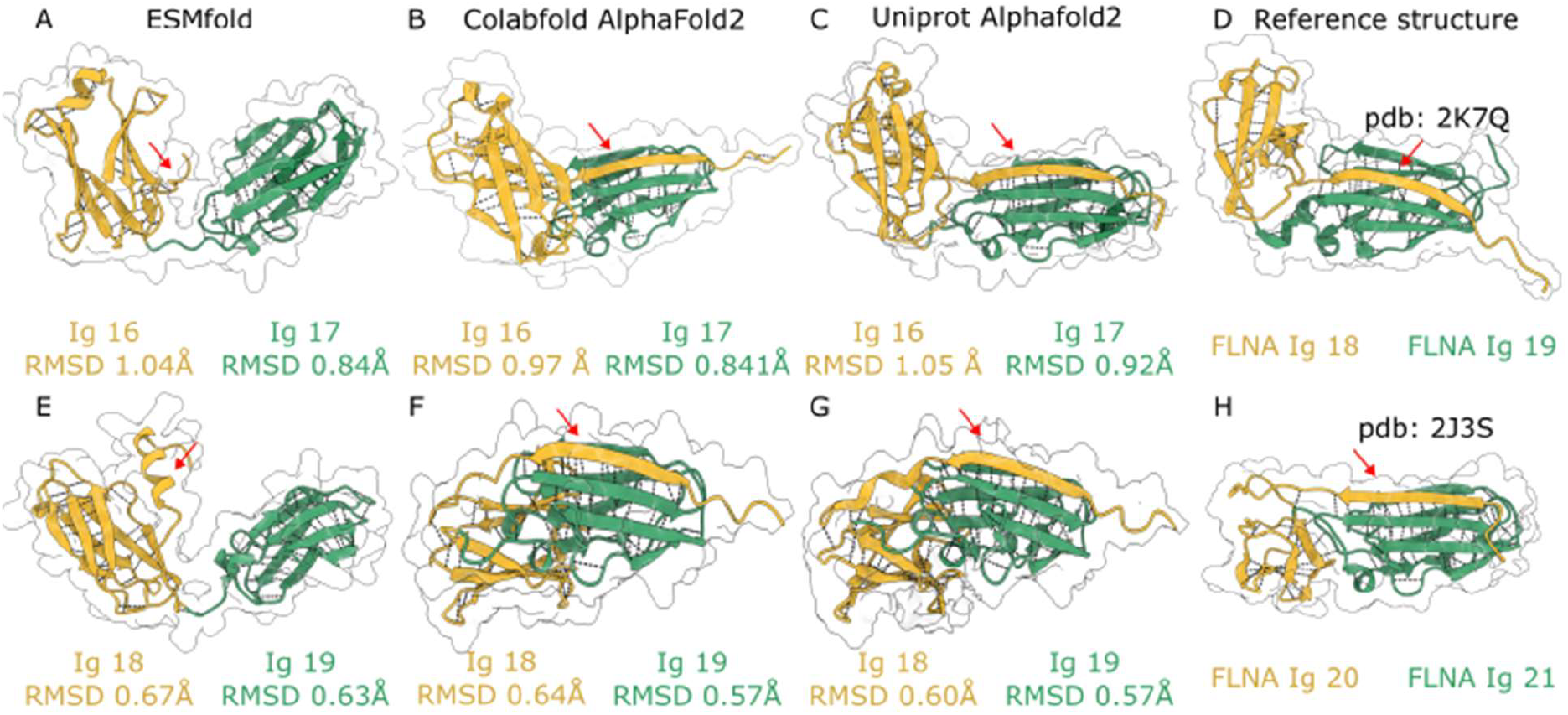
AlphaFold2, but not ESMFold, correctly predicts the closed structure of filamin Ig domains in the MSR. A) Predicted model of Ig16 and Ig17 using ESMFold. B) Predicted model of Ig16 and Ig17 using ColabFold-AlphaFold2. C) Predicted AlphaFold2 model of Ig16 and Ig17 obtained from UniProt (Q9VEN1). D) Crystal structure of Ig18 and Ig19 of human filamin A, homologous to Ig16 and Ig17 of fly filamin, obtained from the Protein Data Bank (PDB: 2K7Q). E) Predicted model of Ig18 and Ig19 using ESMFold. In panels A and E, the masking strand of Ig16 or Ig18 is not attached to Ig17 or Ig19 respectively (red arrows). F) Predicted model of Ig18 and Ig19 using ColabFold. G) Predicted model of Ig18 and Ig19 using AlphaFold 2, obtained from UniProt. H) Crystal structure of Ig20 and Ig21 of human filamin A, homologous to Ig18 and Ig19 of fly filamin, obtained from the Protein Data Bank (PDB: 2J3S). Ig16 and Ig18 are represented in gold, while Ig17 and Ig19 are represented in green. Black dotted lines indicate hydrogen bonds. The red arrow points to the β strand of Ig16 and Ig18 that masks the binding site in Ig17 and Ig19, respectively. The root mean square deviation (RMSD) of each predicted Ig domain compared to the reference structure is provided below each predicted structure.

Filamins are widely expressed proteins involved in many processes and therefore bind to many proteins [1,3]. Traditionally, filamin ligands are identified using various biochemical and biophysical methods such as yeast two-hybrid screens, co-immunoprecipitation assays, and surface plasmon resonance [18,21–23]. However, these methods are often time-consuming and challenging due to the large size of the filamin protein. Recent advances in artificial intelligence and machine learning have led to the development of deep learning-based protein folding models such as AlphaFold2, which can predict the structure of protein-protein complexes with high accuracy [24,25]. In this study, we 1) evaluated the use of AlphaFold2 to identify the binding site of Ig domains 17 and 19 of *Drosophila* filamin; 2) assessed its ability to specifically recognize the open conformation of the MSR; and 3) conducted a biased bioinformatics screen to identify novel mechanically associated filamin-binding partners.

## 2. Methods

### 2.1 Structure Prediction and Model Building

AlphaFold2 model building was conducted using ColabFold v1.5.5 [25], an implementation of AlphaFold2, with the AlphaFold2-multimer-v2 model [24–26]. No template structures were used. For the initial screen, a single model was generated for each pair of sequences. For the follow-up models, three or five models were generated for each pair of sequences with three recycles iteration per model. We obtained the premade AlphaFold2 models from the UniProt Database entry Q9VEN1 [27]. The individual domains were then cropped in ChimeraX [28]. ESMfold model building was performed using the ESM metagenomic Atlas through Fold sequence directly on the esmatlas website [29]. The amino acid sequences used to generate the models are provided (Supporting Information - Text 4). The Root mean square distance (RMSD) was calculated from the generated models to the reference structures from Filamin-A Ig-like domains 18-19 pdb:2K7Q [7] and Ig domains 19 to 21 pdb: 2J3S [14] using the matchmaker command in UCSF ChimeraX v1.9 [28].

### 2.2. Amino acid sequences

We obtained the amino acid sequences from Flybase [30]. We used the longest splice variant, the accession numbers are Cher-PG FBpp0288453, Mys-PD FBpp0309123, Nuak-PD FBpp0293890, HTS-PA FBpp0301112, and Kelch-PB FBpp0080596. For the biased screen, we selected cytoskeleton proteins, myofibril proteins, kinases, proteins involved in protein degradation and filamin binding proteins from the BioGrid database [31]. Then, we obtained the sequences of the longest splice variant using Flybase[30]. The complete list of selected proteins is in the Supporting Information - Text 1. We used the Ig19 domain of Cher-PG as bait. The prey proteins were trimmed to stretches of 200 amino acids and pasted next to Ig19 sequence separated by “:” in fasta file using custom R code (Supporting Information - Text 5).

### 2.3 Data Analysis

Predicted Aligned Error (PAE) quantifies the confidence of AlphaFold2 in the relative positioning of two residues in the predicted structure. It is the expected positional deviation at residue X, measured in Ångströms (Å), assuming the predicted and actual structures are aligned at residue Y. We obtained the PAE plots from the ColabFold v1.5.5 [25].

The modelling results from ColabFold results are a PDB file and a JSON file that contains a matrix of residue paired predicted aligned error (PAE). We used the R library bio3d to open the pdb file in R and make a list of all the contact sites in the model between the two substructures (Ig19 and prey proteins), using a cutoff distance of 5Å [32,33]. Then, we used the R library jsonlite to extract the PAE values at the contact sites specifically [34]. The CPAE values are the average of the PAE values at the contact sites. A general code is in the Supporting Information - Text 6.

To distinguish between the association types occurring between Cher Ig domains and the prey fragments, we generated snapshots of the structures, zooming into the binding site using ChimeraX v1.9 ribbon display[28], which reads the secondary structure from the PDB file. We then counted the number of antiparallel, parallel, and random associations. Both the antiparallel and parallel categories form a β-sheet between the Ig domain and the ligand, but the former uses antiparallel β-strands, while the latter uses parallel β-strands. The random category includes all associations that do not involve a β-sheet. Plots and statistical tests were calculated in R using standalone packages.

## Results

### ColabFold and Alphafold can predict the closed state of filamin Ig domain pairs

Initially, we assessed the performance of two deep learning-based methods in predicting the structure of filamin Ig domains and their corresponding ligands. We utilized ColabFold-AlphaFold2, which combines MMseqs2’s rapid homology search with AlphaFold2-Multimer, and ESMfold, a faster method albeit with lower accuracy than AlphaFold [29]. Specifically, we focused on predicting the structure of Ig domains 16-17 and 18-19 from *Drosophila* typical filamin (Cheerio or Cher). Our rationale behind this comparison was that if these methods could accurately predict the folded conformation of these domain pairs, they would demonstrate the potential to predict novel ligands. Successfully capturing the closed-folded state of these domains would imply their ability to uncover previously unknown binding partners. As a reference structure, we used the crystal structure from Filamin-A Ig domains 18-19 and 19-21, which correspond to 16-17 and 18-19 in *Drosophila* filamin (Fig 1D and H).

When analyzing the structure of Ig16-17, we found that ESMfold successfully predicted two Ig-like domains consisting of 7 β-sheets each (Fig. 1A). However, it failed to predict the β-strand from Ig16 that blocks the binding site in Ig17 (Fig. 1A and D). Similarly, when attempting to predict the structure of Ig domains 18-19 as a pair, ESMfold accurately predicted the individual Ig-like domains but did not capture their interaction (Fig 1E and H). Instead of the expected blocking β-sheet, ESMfold predicted an α-helix (Fig. 1E), which is not consistent with the published structures of verebrate filamins. The pLDDT value of the model was the lowest at the α-helix and highest at the Ig-like domains (Supporting information - Figure 1). Overall, ESMfold cannot predict the closed state of filamin domain pairs and is therefore unlikely to identify mechanobinding partners.

In contrast, ColabFold-AlphaFold2 successfully predicted the presence of a β-strand originating from either Ig16 or Ig18, interacting with the corresponding face of Ig17 or Ig19, respectively (Figure 1B, and E). The pDLLT values at the interface between the two Ig domains were high (Supporting information - Figure 1). Hydrogen bonds at the interdomain interface were also correctly predicted (Fig 1 B, and E). As a control, we used the InterPro-Alphafold2 model from the InterPro database corresponding to the Filamin-PA splice variant (Q9VEN1; Cher-PA). Like our ColabFold-AlphaFold2 results, the InterPro model correctly predicted the closed state of the two domain pairs with high pLDDT values (Figure 1C, F, and Supporting information - Figure 1). To further evaluate the models, we compared them to the crystal structures of FLNA Ig 18-19, which correspond to *Drosophila* Ig domains 16-17, and FLNA Ig 20-21, which correspond to *Drosophila* Ig domains 18-19 (Figure 1D and H). We calculated the root mean square distance (RMSD) between individual Ig domains and the reference structure (Figure 1). ESMfold had the highest RMSD values, ranging from 0.639 Å for Ig 19 to 1.04 Å for Ig16. Then, the InterPro-Alphafold2 model had RMSD values from 0.573 Å for Ig 19 to 1.056 Å for Ig 16. ColabFold-AlphaFold2 provided the best scores with 0.572 Å to 0.974 Å Ig 19 for Ig 16. In all cases, Ig19 has the lowest RMSD values, followed by 18, 17, and 16 (Figure 1). Overall, ColabFold-AlphaFold2 and InterPro-Alphafold2 correctly predicted the closed state in both domain pairs, Ig domains 18-19 are the best predicted. ColabFold-AlphaFold2 models had the lowest RMSD values. Therefore, we restricted our analysis to the ColabFold-AlphaFold2.

### ColabFold-AlphaFold2 correctly predicts known filamin-binding sites

We then tested if we could identify the binding site of filamin on integrin using *Drosophila* proteins. In vertebrates, Ig 21 of Filamin-A, which corresponds to Ig19 of *Drosophila* filamin, binds directly to a single binding site at the cytoplasmic tail of integrin [18,35]. Integrins are transmembrane receptors with a small cytoplasmic tail. Filamin cannot bind to integrin extracellular or transmembrane portions, providing negative controls for predictions. We expect the interaction in *Drosophila* to be like that of vertebrates.

We divided the entire sequence of Myospheroid (Mys, *Drosophila* β subunit of the integrin dimer) into 200 amino acid portions and made ColabFold-AlphaFold2 models of the portions combined with Cher Ig 19. Overall, the predictions were well supported by low Predicted Aligned Error (PAE) values and in all cases the two structures were in direct contact (Figure 2 A-E). The cytoplasmic binding site was identified as an antiparallel β-strand interface (Figure 2E). We noticed that the PAE values at the interfaces were high for the models between Ig-19 and the extracellular portions of Integrin (Fig 2 H-I arrows), suggesting the predicted contact is incorrect. In contrast, the correctly identified binding site had a localized stripe of very low PAE values (Figure 2J arrow). Therefore, we hypothesized that extracting the PAE values at only the contact sites between the two structures would separate the real binding sites formed by antiparallel β-strands from the incorrect ones. We defined the contact sites as residues in the proximity of 5 Å to each other and used the Bio3D package in R to identify them. Then, we extracted the PAE values from those contact sites and referred to these as Contact PAE values (CPAE). The CPAE values of the Integrin to Ig19 models were significantly lower in the correctly predicted cytoplasmic site compared to the extracellular portions (Figure 2K). Therefore, the CPAE allows identifying the correctly predicted binding sites from ColabFold-AlphaFold2 models.

**Figure 2.**
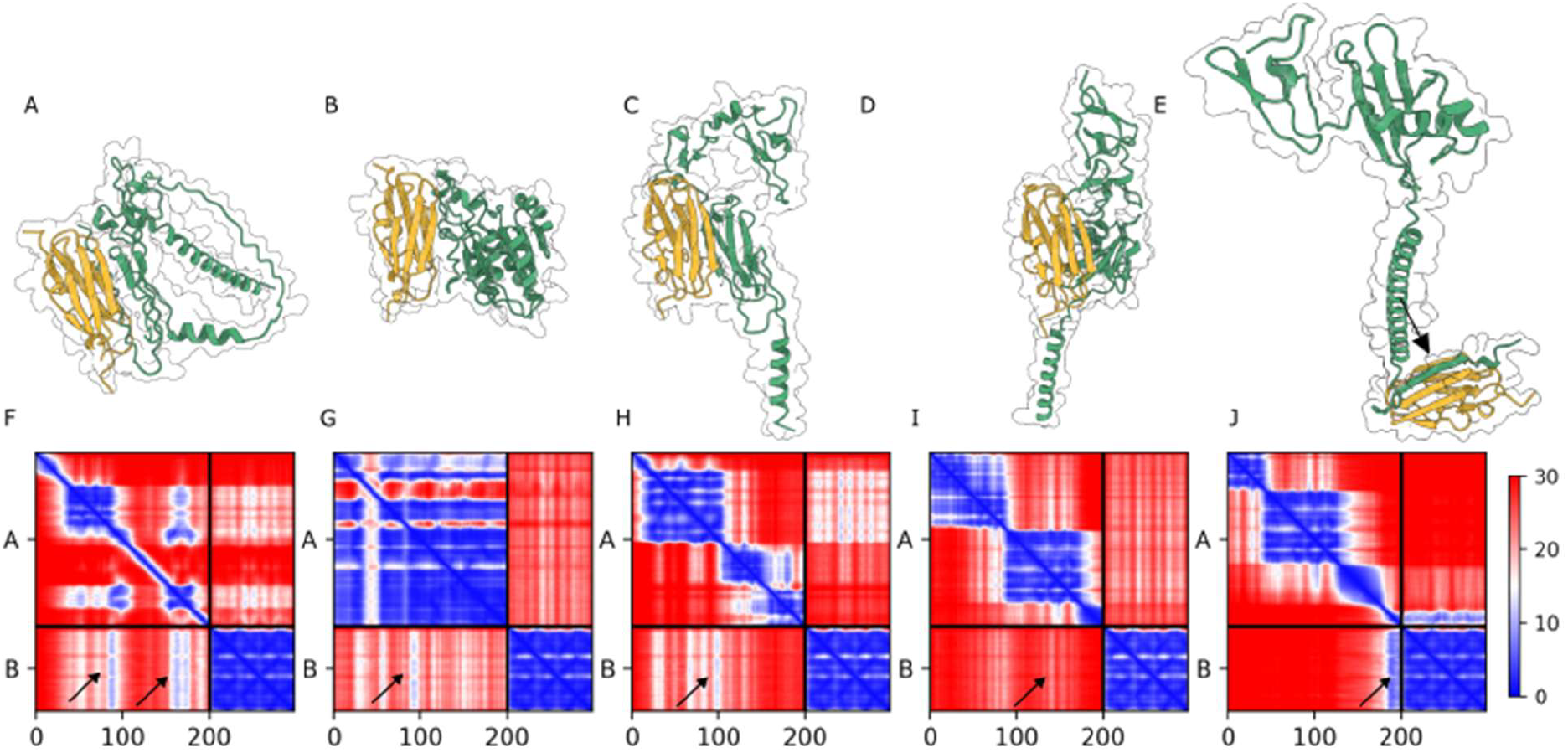
AlphaFold2-multimer accurately identifies the binding site between Drosophila integrin and filamin Ig domain 19. A-E) Predicted structures derived from regions of Mys (green) and Cher Ig19 (gold). Panels A-D contain portions of Mys that are in the extracellular space and therefore should not be associated with filamin. Panel E contains an extended (α helix corresponding to the single transmembrane pass and a short intracellular portion, containing the correct binding site (arrow). F-J) PAE plots depicting the predicted structures. The color scale on the right indicates confidence levels, with dark blue regions representing high-confidence binding sites. A and B denote chains in the PAE plots, and the intersection between them represents the scores between the chains. Dark blue regions at the intersections correspond to potential contact sites, while regions with higher PAE values indicate non-contact regions.

Then we tested the specificity of filamin’s Ig domains on recognizing integrin’s cytoplasmic tail. We used the Ig domains from the mechanosensitive region of Cher (Ig 14-19), which correspond to human filamins Ig domains 16-21. We made 5 models for each Ig domain in combination with the cytoplasmic tail of Mys. Ig domains 15, 17, 18, and 19 were always predicted as antiparallel β-strands, the rest were never predicted as antiparallel β-strands (Figure 3A). Then, we calculated the CPAE values of the models. As positive control, we made a ColabFold-AlphaFold2 model of Human Filamin-A Ig21 with the Human integrin beta7 cytoplasmic tail, for which crystal structures are available (2BRQ and 2JF1 [35,36]). We noticed that the CPAE values of the models were very low (<10) when the antiparallel binding site was modelled and high (>10) when it was not (Figure 3B). Ig15, 17, and 19 had the lowest CPAE values (Figure 3B). Ig18 was modelled forming an antiparallel β-strand with the integrin tail but with higher CPAE values. Since Ig18 is the blocking domain, it is unlikely to bind to the integrin tail.

**Figure 3.**
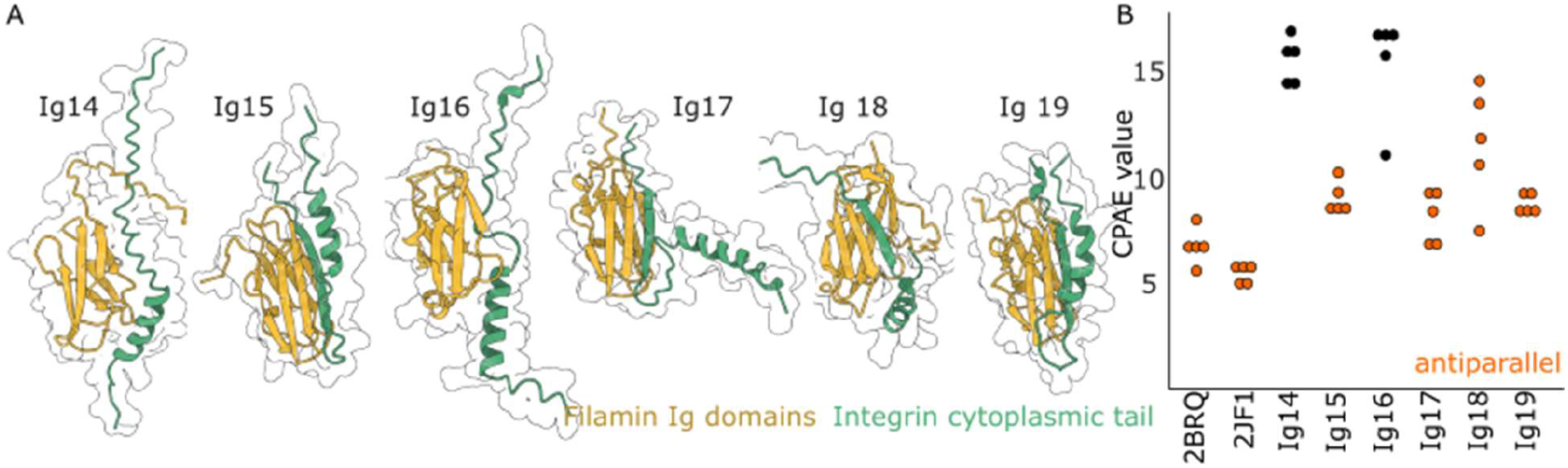
AlphaFold2 recognizes the integrin-filamin correct binding sites. (A) Representative AlphaFold2 models of the Mys integrin cytoplasmic tail from with the six Ig domains that compose Cher MSR. (B) Plot showing the CPAE scores for each Ig domain of Cher with the cytoplasmic tail of Mys. As controls, we used the structures of human filamin with integrin tail (2BRQ and 2F1).

### Identification of binding sites in Kelch, Nuak, and Hts

We then investigated the association between three well-supported filamin interactors for which the precise binding sites remain unknown, Kelch, Nuak[23], and Hts[37]. We began with Kelch, a Cullin3-RING ubiquitin E3 ligase component involved in substrate targeting [38–41]. We obtained the sequence from the Kelch-PB splice variant and divided it into 200 amino acid regions. We performed structure predictions by combining these regions with all 22 Ig domains of Cher. Most of the predictions had CPAE values above 10 and seven predictions below had CPAE values below 10 (Figure 4A). Among these, six exhibited an expected antiparallel β-strand on β-strand contact. Specifically, we identified three potential binding sites in Kelch: Binding site 1 (ASSFFSCLH) interacted with Ig17, Binding site 2 (AVGGAVA) interacted with Ig15, Ig17, and Ig19, and binding site 3 (VGHIRLNA) interacted with Ig8. Notably, all the identified binding sites in Kelch were located within the C-terminus region, which is known to mediate substrate binding for protein degradation [40].

**Figure 4.**
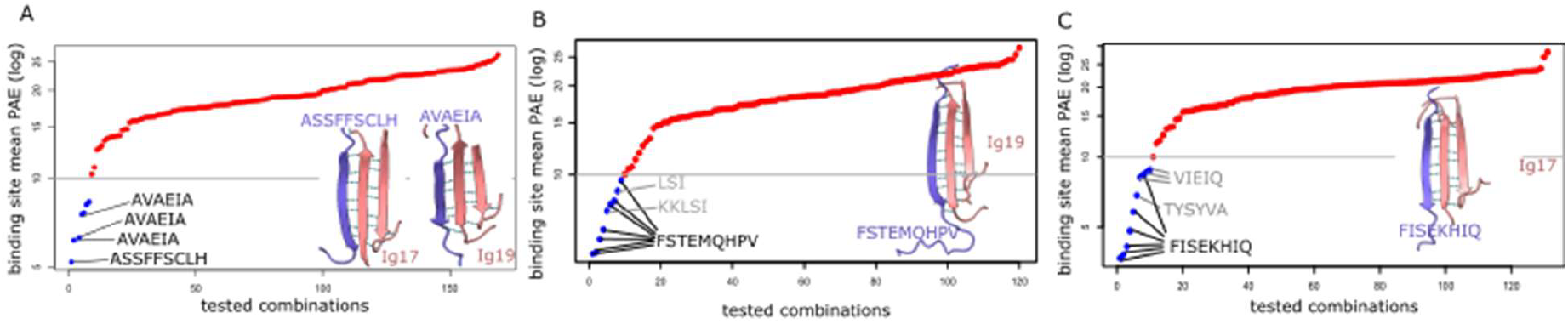
Predicted binding sites for three well-known filamin interactors. A) Plots of the Contact Predicted Aligned Error (CPAE) values at the contact sites between filamin Ig domains and portions of Kelch. B) Plot of PAE values evaluating the contact sites between portions of NUAK and filamin Ig domains. C) Plot of PAE values assessing the contact sites between portions of Hts and filamin Ig domains. The binding sites of the predicted models are inside the plot. The Y-axis are on a log scale. The tested interactions are sorted based on their PAE scores.

We then proceeded to investigate the interaction between the protein Nuak and Cher. Nuak is a serine/threonine kinase involved in muscle autophagy regulation [23,42,43]. Again, we paired portions of 200 amino acids from Nuak with individual Cher Ig domains. The interactions were evaluated based on their CPAE scores. From these evaluations, we identified eight interactions with the sequence FSTEMQHPV as the putative binding site in Nuak (Figure 4B). Cher Ig15, Ig17, and Ig19 had the lowest CPAE scores in these interactions (Figure 4B). Two other β-strand forming sites containing the residues KKLSI and LSI were also recovered, but they are too short to constitute real binding sites. Nuak is predominantly a disordered protein, except for a small kinase domain located at the N-terminus (residues 71-321 in Nuak-PG). The predicted binding site in Nuak corresponds to residues 843-851, which are in the disordered region.

Finally, we investigated the interaction between Cher and Hu-li tai shao (Hts), an actin-binding protein that interacts with Cher at the ring canals [37,44–46]. Again, we paired portions of 200 amino acids from Hts with individual Cher Ig domains. We used the longest Hts splice variant Hts-PA. We recovered 10 models with CPAE values lower than 10 all of them forming β-strand at the interface with Cher domains (Figure 4C). Six of these models identified the same binding site in Hts corresponding to the residues 835-842 FISEKHIQ and five of these had the lowest CPAE values (Figure 4C). Two other binding sites, TYSYVA and VIEIQ, had slightly higher CPAE values. At the ring canal, Hts is cleaved to a smaller form called Hts-RC (which stands for Ring Canal). The RC form is produced through proteolytic cleavage at position 692 and corresponds to the C-terminal portion of the cleavage. Therefore, the interaction must occur somewhere between residues 692 and 1156. The binding site FISEKHIQ which had the lowest CPAE values falls into the Hts-RC form, The site VIEIQ is located at residues 639-642 just before the start of Hts-RC and therefore may represent a false positive. The binding site TYSYVA corresponds to residues 1082 to 1087 which also corresponds to the RC form.

### Screen for novel filamin-binding proteins

Once we established that AlphaFold2 can accurately identify the binding sites of known filamin-binding proteins, we decided to use it to uncover novel filamin-binding proteins. To reduce the computational load, we restricted our search to Ig19. Since we are interested in the mechanosignaling function of filamin at the myofibrils[22,47–49], we limited the screened candidates to proteins that are myofibril components, cytoskeleton proteins, listed as possible filamin interactors based on the BioGRID project, kinases and proteins related to protein degradation [30,31]. We obtained a list of ∼1000 proteins (Supporting Information - Text 1). We downloaded their amino acid sequences from FlyBase and split them into regions of 200 amino acids and paired them with Ig19 (Figure 5A). In total, we generated 5247 models corresponding to 946 proteins. From these, we obtained the CPAE values and sorted them accordingly. 132 had a CPAE value below 5, corresponding to 84 proteins. 834 models had CPAE values below 10, corresponding to 365 proteins (Figure 5B). Known interactions such as Kelch, Sls, Hts, Nuak, and Mys were recovered, confirming the approach (Supporting information - Text 2 and 3). We identified three types of interactions between the candidate segments and Ig19. The first type was an antiparallel β-sheet formed between Ig19 and a stretch of amino acids in the other protein. The second type was a parallel β-sheet. The third type involved the random juxtaposition of the two structures without a defined pattern (Figure 5C).

**Figure 5.**
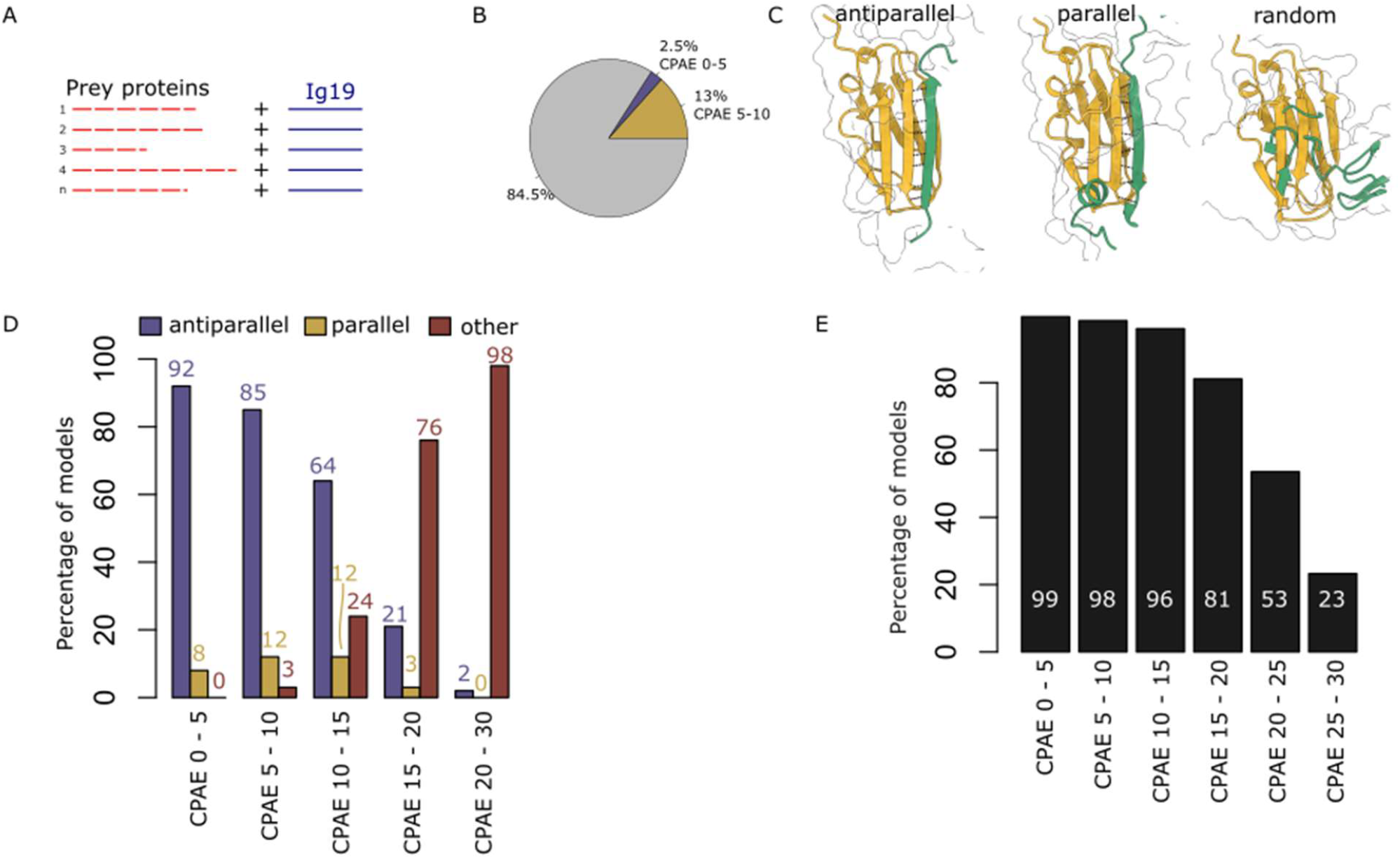
Details of the screen for Ig19 binding targets. (A) Prey proteins are segmented into smaller fragments of 200 amino acids, which are then paired with the Ig19 domain of filamin and modelled through AlphaFold2-ColaFold. (B) Pie chart showing the efficiency of the screen at thresholds of CPAE<5 and CPAE<10. (C) Predicted structures of prey protein fragments (green) and filamin Ig19 (gold) show three types of associations: antiparallel binding of the prey protein fragment with filamin Ig19, parallel binding, and random, which corresponds to the random juxtaposition of the two structures in space. (D) Bar plot showing the percentage of predicted models in each conformation category across bins of CPAE values. As CPAE values increase, the percentage of antiparallel binding conformations decreases, while the percentage of random conformations increases. Significance was calculated using a proportion test followed by a Bonferroni correction for multiple comparisons. The p-values are antiparallel p = 2e-51, parallel (not significant), and random p = 9e-68. (E) Percentage of models in which contacts occur at positions 38–45 of Ig19.

We observed that 92% of models with CPAE < 5 correspond to an antiparallel β-sheet, with this percentage decreasing sharply as CPAE values increase (Figure 5D). In contrast, the number of parallel β-sheets remained relatively unchanged as CPAE values increased (Figure 5D), suggesting that parallel β-sheets may be artifacts. Lastly, models with a random juxtaposition of the two structures dominated high CPAE models but were almost absent in low CPAE models (Figure 5D), implying that these structures do not represent genuine interactions.

Most models showed contacts at positions 38–45 (Figure 5E), corresponding to the well-established binding site of Ig19 (Figure 5E). Among models with CPAE < 5, 98% had contacts at the correct sites. This percentage remained stable for models with CPAE values up to 15 but declined for models with higher CPAE values. We also tested for biases in the positions of the residues in the prey proteins, as no preferred positions were expected. We did not observe any strongly enriched regions in the dataset (Supporting information - Figure 2), suggesting that the model does not exhibit a strong positional bias. Finally, we looked for conserved residues in models with CPAE < 5 using a multiple-sequence alignment tool. However, we did not identify a clear motif or pattern (Supporting information - Figure 3).

### Differences between Ig19 and Ig17

Ig17 and Ig19 are very similar in structure and function. We therefore asked whether the screen we performed against Ig19 would enrich for Ig19 ligands over Ig17 binders. To test this, we generated new models for the 255 candidates with CPAE values less than 5 from the Ig19 screen, using either Ig17 or Ig19 as targets. For Ig19, the models showed consistently low average CPAE values, demonstrating the reproducibility of the screening approach (Figure 6A), although a few models exhibited higher CPAE values. Similarly, for Ig17, the models had lower CPAE values compared to random models from the screen (Figure 6B). This suggests that screening for Ig19 indirectly selects for Ig17 binders. Overall, the mean CPAE value of all models was lower for Ig19 than for Ig17 (Figure 6C). Next, we examined specific interactions with each Ig domain. The models were divided into three groups: those with no change in CPAE value, those with higher CPAE values in Ig19, and those with lower CPAE values in Ig19 (Figure 6D). Most models showed no change in CPAE between the Ig domains, suggesting that they recognize both domains. Of the 255 models, 34 had lower CPAE values in Ig19, and 12 of these had CPAE values below 5, indicating specific recognition of Ig19. For example, we identified a binding site in the protein Linear Ubiquitin E3 Ligase (LUBEL) that Ig19 recognizes but not Ig17 (Figure H). Additionally, we found a binding site in NUAK that is recognized by both Ig domains (Figure 6E). In contrast, only 2 models had lower CPAE values in Ig17, and only one of these had a CPAE value below 5. A closer inspection of the model revealed an uncommon interaction between Slik and the Ig domains, characterized by the presence of two β-strands instead of the typical single strand (Figure 6K).

**Figure 6.**
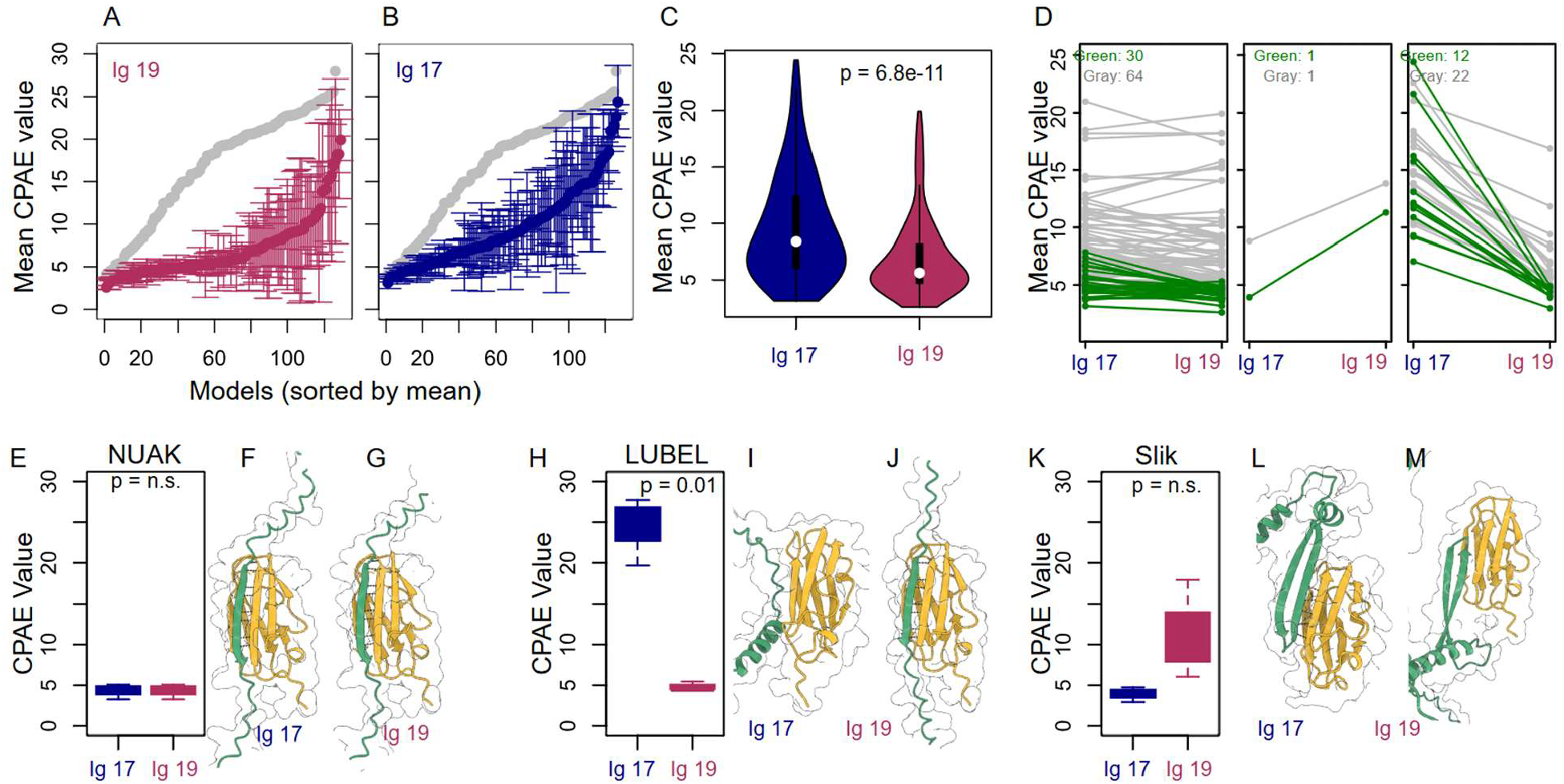
Models selected for Ig19 are either specific to Ig19 or common between Ig17 and Ig19. (A) A plot of CPAE values from the top 130 hits against Ig17, sorted by mean CPAE value; standard deviation is shown. (B) A plot of CPAE values from the top hits against Ig19; standard deviation is shown. The gray line in panels A and B represents the baseline obtained from sampling 130 random models from the original screen. The values in A and B are consistently lower than the baseline. (C) Violin plot of the mean CPAE values for the top hits against Ig17 and Ig19, showing that Ig19 values are lower. A paired t-test was used to calculate the p-value at a 95% confidence level. (D) Paired slope plots comparing values for the top hits against Ig17 and Ig19, categorized into three groups: those with no change between Ig domains (left), those lower in Ig17 (middle), and those lower in Ig19 (right). Models with at least one mean value below 5 CPAE are highlighted, while the rest are shown in gray. Many models are common between Ig domains (left plot) and lower in Ig19 (right plot), with only one model showing lower values in Ig17. (E) Boxplot of CPAE values for a portion of the NUAK protein paired against Ig17 and Ig19. (F) Model of NUAK paired with Ig17. (G) Model of NUAK paired with Ig19. In both models, an antiparallel beta-sheet forms between the structures. (H) Boxplot of CPAE values of models of a portion of the LUBEL protein paired with Ig17 and Ig19. (I) Model of LUBEL paired with Ig17. (J) Model of LUBEL paired with Ig19. An antiparallel beta-sheet forms with Ig19 but not with Ig17. (K) Boxplot of CPAE values for models of Ig17 and Ig19 paired with a portion of the Slik protein. (L, M) Models of Ig17 and Ig19 with a portion of Slik, show that antiparallel beta-sheets do not form in either model. In C, E-K, statistical significance was calculated using Welch Two Sample t-tests, n.s. = not significant.

### High-confidence ligands do not recognize the closed MSR conformation

Finally, we asked whether the top-ranked binding regions specifically recognize the open MSR conformation. To test this, we created new models using the binding regions from CPAE < 5 models, now in combination with Ig17 or Ig19, which represent the open conformation, and in combination with Ig16-Ig17 and Ig18-Ig19, which are typically modelled in closed conformations. Overall, the CPAE of all the new models was significantly lower in the models that included Ig17 or Ig19 alone compared to those that included the blocking Ig domains, i.e., Ig16-Ig17 and Ig18-Ig19 (Figure 7A). Similarly, the percentage of models in which an antiparallel β-strand was modelled as the interaction was the majority when using Ig17 or Ig19 alone but was very low when using Ig16-Ig17 and almost absent when using Ig18-Ig19 (Figure 7B). As an example, we chose one ligand that forms antiparallel β-strands with Ig17 and Ig19 found in the protein Sterile20-like kinase (Slik). It has low CPAE values with Ig17 or Ig19 but high values with Ig16-Ig17 or Ig18-Ig19 (Figure 7C). The residues VTTAIEVAI in Slik bind to Ig19 at the spot where the blocking strand from Ig18 sits (Figure 7E and E). However, when modelled together with Ig18-Ig19, the blocking strand from Ig18 blocks the interaction between the VTTAIEVAI residues and Ig19 (Figure 7F). Overall, the binding regions confidently recognize the open conformation of either Ig17 or Ig19 but do not confidently recognize the closed conformations.

**Figure 7.**
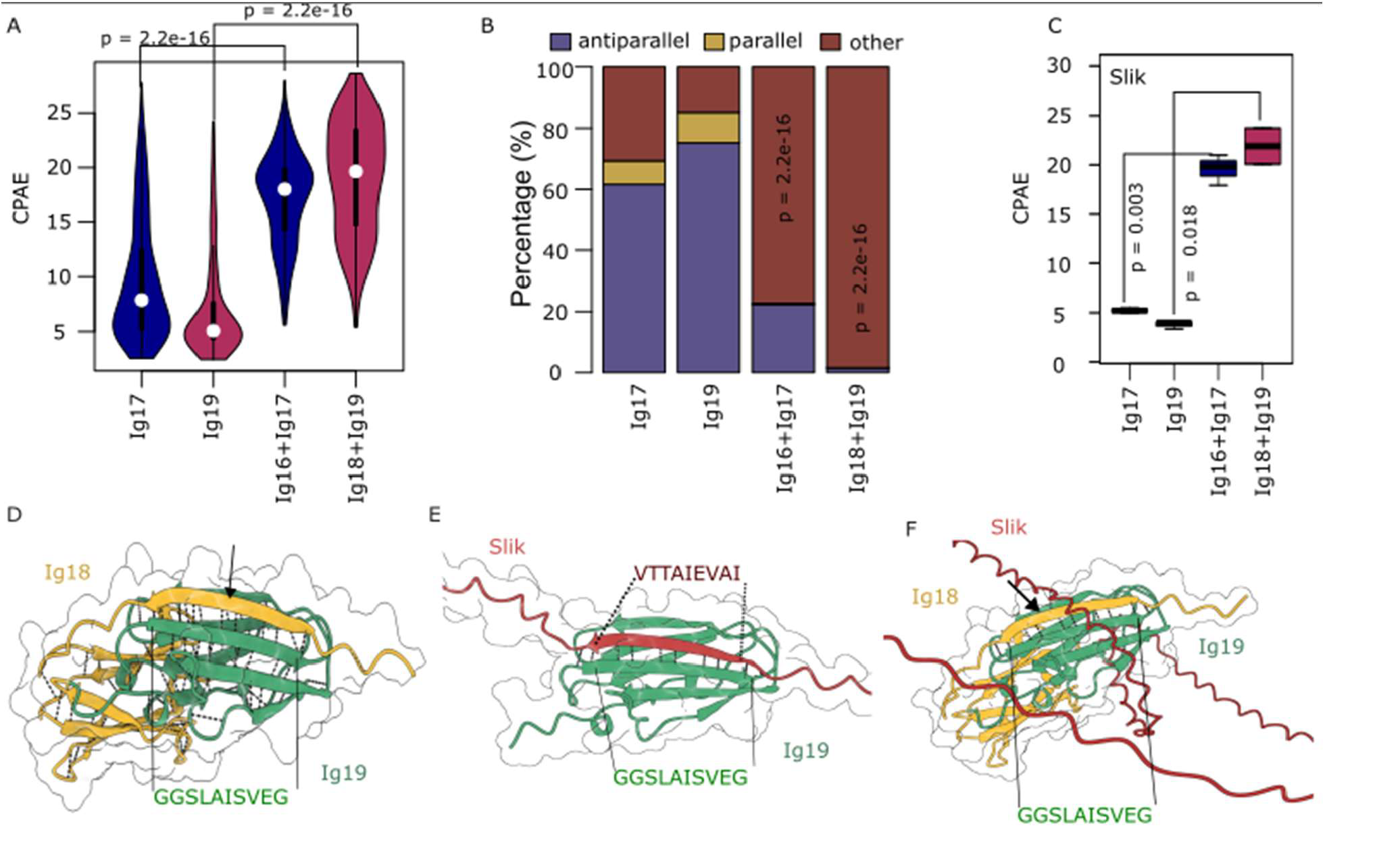
High confidence ligands do not recognize the closed version of the MSR. (A) Violin plots of CPAE values of high confidence ligands obtained from the screen retested with Ig17, Ig19, Ig16-Ig17, and Ig18-Ig19. Ig 17 and Ig19 by themselves represent the open conformations, while Ig16-Ig17 and Ig18-Ig19 represent the closed conformations. Statistical significance was calculated using Welch Two Sample t-tests (B) Stacked plot shows the percentage of predicted models in the three contact categories. Statistical significance was calculated using a Pearson’s Chi-squared test (C) Plot of CPAE values of a region in Slik which oddly forms antiparallel B-strands with Ig16-Ig17. Statistical significance was calculated using Welch Two Sample t-tests. D) Model of Ig18-Ig19 showing the binding site in Ig19 and blocking strand (arrow). E) Model of the interaction between Slik and Ig19. (F) Model of the interaction between Slik and Ig18-Ig19, note the blocking strand in Ig18 prevent Slik binding (arrow).

## Discussion

In this study, we evaluated an AlphaFold2 approach to identify proteins that bind to Ig19, one of the mechanically induced binding sites in *Drosophila* filamin, using ColabFold notebooks [25]. The following evidence supports the method: 1) AlphaFold2 can accurately predict binding events for the cytoplasmic tails of integrins at the correct binding site. 2) AlphaFold2 identified the binding sites for Nuak, Hts, and Kelch—three proteins known to interact with filamin but lacking clearly identified binding sites. In all cases, a high-confidence region was identified, which aligns well with previous genetic and biochemical experiments [22,23,39,45]. 3) A biased screen for Ig19 ligands uncovered mostly structures forming antiparallel β-strands with Ig19, which is the standard mechanism by which filamin Ig domains recognize their targets [8,13–18]. 4) The identified ligands do not recognize the closed conformation.

Low CPAE values and antiparallel β-strands interphases are strong indicators of potential binding interactions. It is well established that the interphase between Ig17 or Ig19 and their ligands forms an antiparallel β-strand interphase [1,3]. Most of the low CPAE models we recovered had antiparallel β-strand interphases, but some did not. Those with different interphases may represent false positives. We are confident in the approach since β-Strands are very common motif for Protein–Protein Interfaces and are well represented in AlphaFold2 training data [50].

Filamins are multitarget scaffold proteins, and some of their ligands are sensitive to the state of the filamin MSR[1,3]. *In-vivo* approaches have been used to identify proteins that are in a complex with filamin in *Drosophila* [21,37]. However, since filamin ligands are tissue-dependent, these methods cannot identify ligands in tissues other than those in which the experiment was performed. While *in-vitro* methods, such as yeast two-hybrid screens, can, in principle, test all proteins in an organism, they suffer from high rates of false positives [51]. Another limitation of these approaches is their inability to resolve specific binding sites. In contrast, AlphaFold2 screens can provide insights into protein-protein interactions at a much higher resolution. When combined with tissue-specific gene expression databases, one could select only those proteins expressed in a tissue of interest, significantly reducing the computational costs of the screen. Thankfully, there are many RNA-seq databases available at the tissue or single-cell level for *Drosophila* [30,52–54].

## Conclusion

Alphafold2 allows the rapid screening of filamin ligands, especially the ones that bind at the masked binding sites of Ig 17 and 19 of *Drosophila* filamin.

## Acknowledgments

Our research is funded by grants to NGM from NSERC (RGPIN/02984 -2022 and DGECR/00166 - 2022), the Canada Foundation for Innovation and Research Nova Scotia (CFI 43947 2023-2795).

## References

1. Razinia Z, Mäkelä T, Ylänne J, Calderwood DA. Filamins in mechanosensing and signaling. Annual Review of Biophysics. 2012. doi:10.1146/annurev-biophys-050511-102252

2. Nakamura F, Osborn TM, Hartemink CA, Hartwig JH, Stossel TP. Structural basis of filamin A functions. Journal of Cell Biology. 2007;179. doi:10.1083/jcb.200707073

3. Mulder T, Johnson J, González-Morales N. The filamins of Drosophila. Genome. 2025;68: 1–11. doi:10.1139/gen-2024-0159

4. Huelsmann S, Rintanen N, Sethi R, Brown NH, Ylänne J. Evidence for the mechanosensor function of filamin in tissue development. Sci Rep. 2016;6. doi:10.1038/srep32798

5. Rognoni L, Stigler J, Pelz B, Ylänne J, Rief M. Dynamic force sensing of filamin revealed in single-molecule experiments. Proc Natl Acad Sci U S A. 2012;109. doi:10.1073/pnas.1211274109

6. Lad Y, Kiema T, Jiang P, Pentikäinen OT, Coles CH, Campbell ID, et al. Structure of three tandem filamin domains reveals auto-inhibition of ligand binding. EMBO Journal. 2007;26. doi:10.1038/sj.emboj.7601827

7. Heikkinen OK, Ruskamo S, Konarev P V., Svergun DI, Iivanainen T, Heikkinen SM, et al. Atomic structures of two novel immunoglobulin-like domain pairs in the actin cross-linking protein filamin. Journal of Biological Chemistry. 2009;284. doi:10.1074/jbc.M109.019661

8. Pentikäinen U, Ylänne J. The Regulation Mechanism for the Auto-Inhibition of Binding of Human Filamin A to Integrin. J Mol Biol. 2009;393. doi:10.1016/j.jmb.2009.08.035

9. Deng Y, Yan J. Force-Dependent Structural Changes of Filamin C Rod Domains Regulated by Filamin C Dimer. J Am Chem Soc. 2023;145. doi:10.1021/jacs.3c02303

10. Pudas R, Kiema TR, Butler PJG, Stewart M, Ylänne J. Structural basis for vertebrate filamin dimerization. Structure. 2005;13. doi:10.1016/j.str.2004.10.014

11. Seo MD, Seok SH, Im H, Kwon AR, Lee SJ, Kim HR, et al. Crystal structure of the dimerization domain of human filamin A. Proteins: Structure, Function and Bioinformatics. 2009;75. doi:10.1002/prot.22336

12. Sokol NS, Cooley L. Drosophila Filamin encoded by the cheerio locus is a component of ovarian ring canals. Current Biology. 1999;9. doi:10.1016/S0960-9822(99)80502-8

13. Ithychanda SS, Das M, Ma YQ, Ding K, Wang X, Gupta S, et al. Migfilin, a molecular switch in regulation of integrin activation. Journal of Biological Chemistry. 2009;284. doi:10.1074/jbc.M807719200

14. Lad Y, Jiang P, Ruskamo S, Harburger DS, Ylänne J, Campbell ID, et al. Structural basis of the migfilin-filamin interaction and competition with integrin β tails. Journal of Biological Chemistry. 2008;283. doi:10.1074/jbc.M802592200

15. Nakamura F, Heikkinen O, Pentikäinen OT, Osborn TM, Kasza KE, Weitz DA, et al. Molecular basis of filamin A-FilGAP interaction and its impairment in congenital disorders associated with Filamin A mutations. PLoS One. 2009;4. doi:10.1371/journal.pone.0004928

16. Ehrlicher AJ, Nakamura F, Hartwig JH, Weitz DA, Stossel TP. Mechanical strain in actin networks regulates FilGAP and integrin binding to filamin A. Nature. 2011;478. doi:10.1038/nature10430

17. Smith L, Page RC, Xu Z, Kohli E, Litman P, Nix JC, et al. Biochemical basis of the interaction between cystic fibrosis transmembrane conductance regulator and immunoglobulin-like repeats of filamin. Journal of Biological Chemistry. 2010;285. doi:10.1074/jbc.M109.080911

18. Liu J, Das M, Yang J, Ithychanda SS, Yakubenko VP, Plow EF, et al. Structural mechanism of integrin inactivation by filamin. Nat Struct Mol Biol. 2015;22. doi:10.1038/nsmb.2999

19. Kesner BA, Ding F, Temple BR, Dokholyan N V. N-terminal strands of filamin Ig domains act as a conformational switch under biological forces. Proteins: Structure, Function and Bioinformatics. 2010;78. doi:10.1002/prot.22479

20. Rognoni L, Möst T, Žoldák G, Rief M. Force-dependent isomerization kinetics of a highly conserved proline switch modulates the mechanosensing region of filamin. Proc Natl Acad Sci U S A. 2014;111. doi:10.1073/pnas.1319448111

21. Külshammer E, Kilinc M, Csordás G, Bresser T, Nolte H, Uhlirova M. The mechanosensor Filamin A/Cheerio promotes tumourigenesis via specific interactions with components of the cell cortex. FEBS Journal. 2022;289. doi:10.1111/febs.16408

22. González-Morales N, Holenka TK, Schöck F. Filamin actin-binding and titin-binding fulfill distinct functions in Z-disc cohesion. PLoS Genet. 2017;13. doi:10.1371/journal.pgen.1006880

23. Brooks D, Naeem F, Stetsiv M, Goetting SC, Bawa S, Green N, et al. Drosophila NUAK functions with Starvin/BAG3 in autophagic protein turnover. PLoS Genet. 2020;16. doi:10.1371/journal.pgen.1008700

24. Evans R, O’Neill M, Pritzel A, Antropova N, Senior A, Green T, et al. Protein complex prediction with AlphaFold-Multimer. bioRxiv. 2022.

25. Mirdita M, Schütze K, Moriwaki Y, Heo L, Ovchinnikov S, Steinegger M. ColabFold: making protein folding accessible to all. Nat Methods. 2022;19. doi:10.1038/s41592-022-01488-1

26. Jumper J, Evans R, Pritzel A, Green T, Figurnov M, Ronneberger O, et al. Highly accurate protein structure prediction with AlphaFold. Nature. 2021;596. doi:10.1038/s41586-021-03819-2

27. Varadi M, Anyango S, Deshpande M, Nair S, Natassia C, Yordanova G, et al. AlphaFold Protein Structure Database: Massively expanding the structural coverage of protein-sequence space with high-accuracy models. Nucleic Acids Res. 2022;50. doi:10.1093/nar/gkab1061

28. Meng EC, Goddard TD, Pettersen EF, Couch GS, Pearson ZJ, Morris JH, et al. UCSF ChimeraX: Tools for structure building and analysis. Protein Science. 2023;32. doi:10.1002/pro.4792

29. Lin Z, Akin H, Rao R, Hie B, Zhu Z, Lu W, et al. Evolutionary-scale prediction of atomic-level protein structure with a language model. Science (1979). 2023;379. doi:10.1126/science.ade2574

30. Gramates LS, Agapite J, Attrill H, Calvi BR, Crosby MA, dos Santos G, et al. FlyBase: a guided tour of highlighted features. Genetics. 2022;220. doi:10.1093/genetics/iyac035

31. Oughtred R, Rust J, Chang C, Breitkreutz BJ, Stark C, Willems A, et al. The BioGRID database: A comprehensive biomedical resource of curated protein, genetic, and chemical interactions. Protein Science. 2021;30. doi:10.1002/pro.3978

32. Grant BJ, Skjærven L, Yao XQ. The Bio3D packages for structural bioinformatics. Protein Science. 2021;30. doi:10.1002/pro.3923

33. Grant BJ, Rodrigues APC, ElSawy KM, McCammon JA, Caves LSD. Bio3d: An R package for the comparative analysis of protein structures. Bioinformatics. 2006;22. doi:10.1093/bioinformatics/btl461

34. Ooms J. The jsonlite Package: A Practical and Consistent Mapping Between JSON Data and R Objects. 2014. Available: http://arxiv.org/abs/1403.2805

35. Kiema T, Lad Y, Jiang P, Oxley CL, Baldassarre M, Wegener KL, et al. The molecular basis of filamin binding to integrins and competition with talin. Mol Cell. 2006;21. doi:10.1016/j.molcel.2006.01.011

36. Takala H, Nurminen E, Nurmi SM, Aatonen M, Strandin T, Takatalo M, et al. 22 integrin phosphorylation on Thr758 acts as a molecular switch to regulate 14-3-3 and filamin binding. Blood. 2008;112. doi:10.1182/blood-2007-12-127795

37. Mannix KM, Starble RM, Kaufman RS, Cooley L. Proximity labeling reveals novel interactomes in live Drosophila tissue. Development (Cambridge). 2019. doi:10.1242/DEV.176644

38. Kelso RJ, Hudson AM, Cooley L. Drosophila Kelch regulates actin organization via Src64-dependent tyrosine phosphorylation. Journal of Cell Biology. 2002;156. doi:10.1083/jcb.200110063

39. Hudson AM, Mannix KM, Gerdes JA, Kottemann MC, Cooley L. Targeted substrate degradation by kelch controls the actin cytoskeleton during ring canal expansion. Development (Cambridge). 2019;146. doi:10.1242/dev.169219

40. Hudson AM, Cooley L. Drosophila Kelch functions with Cullin-3 to organize the ring canal actin cytoskeleton. Journal of Cell Biology. 2010;188. doi:10.1083/jcb.200909017

41. Hudson AM, Mannix KM, Cooley L. Actin cytoskeletal organization in drosophila germline ring canals depends on kelch function in a Cullin-RING E3 ligase. Genetics. 2015;201. doi:10.1534/genetics.115.181289

42. Zhao Z, Brooks D, Guo Y, Geisbrecht ER. Identification of CryAB as a target of NUAK kinase activity in Drosophila muscle tissue. Genetics. 2023;225. doi:10.1093/genetics/iyad167

43. Brooks D, Bawa S, Bontrager A, Stetsiv M, Guo Y, Geisbrecht ER. Independent pathways control muscle tissue size and sarcomere remodeling. Dev Biol. 2022;490. doi:10.1016/j.ydbio.2022.06.014

44. Yue L, Spradling AC. hu-li tai shao, a gene required for ring canal formation during Drosophila oogenesis, encodes a homolog of adducin. Genes Dev. 1992;6. doi:10.1101/gad.6.12b.2443

45. Gerdes JA, Mannix KM, Hudson AM, Cooley L. HtsRc-mediated accumulation of f-actin regulates ring canal size during drosophila melanogaster oogenesis. Genetics. 2020;216. doi:10.1534/genetics.120.303629

46. Petrella LN, Smith-Leiker T, Cooley L. The Ovhts polyprotein is cleaved to produce fusome and ring canal proteins required for Drosophila oogenesis. Development. 2007;134. doi:10.1242/dev.02766

47. Schöck F, González-Morales N. The insect perspective on Z-disc structure and biology. Journal of Cell Science. 2022. doi:10.1242/jcs.260179

48. González-Morales N, Schöck F. Commentary: Nanoscopy reveals the layered organization of the sarcomeric H-zone and I-band complexes. Frontiers in Cell and Developmental Biology. 2020. doi:10.3389/fcell.2020.00074

49. Fisher LAB, Carriquí-Madroñal B, Mulder T, Huelsmann S, Schöck F, González-Morales N. Filamin protects myofibrils from contractile damage through changes in its mechanosensory region. PLoS Genet. 2024;20: e1011101. doi:10.1371/journal.pgen.1011101

50. Watkins AM, Arora PS. Anatomy of β-strands at protein-protein interfaces. ACS Chem Biol. 2014;9. doi:10.1021/cb500241y

51. Formstecher E, Aresta S, Collura V, Hamburger A, Meil A, Trehin A, et al. Protein interaction mapping: A Drosophila case study. Genome Res. 2005;15. doi:10.1101/gr.2659105

52. Brown JB, Boley N, Eisman R, May GE, Stoiber MH, Duff MO, et al. Diversity and dynamics of the Drosophila transcriptome. Nature. 2014;512. doi:10.1038/nature12962

53. Chintapalli VR, Wang J, Dow JAT. Using FlyAtlas to identify better Drosophila melanogaster models of human disease. Nature Genetics. 2007. doi:10.1038/ng2049

54. Li H, Janssens J, de Waegeneer M, Kolluru SS, Davie K, Gardeux V, et al. Fly Cell Atlas: A single-nucleus transcriptomic atlas of the adult fruit fly. Science (1979). 2022;375. doi:10.1126/science.abk2432

