## Supporting Information for "Computational Screening of Filamin Mechanical Binding Proteins using AlphaFold2"

**Supplementary text 1. List of selected proteins included in the biased screen**

14-3-3zeta, Abl, Acaa, Acbp1, AcCoAS, Ack, Ack-like, Act57B, Act79B, Act88F, Actn, Adck1, Adh, Adk1, Adk2, Adk3, Adsl, Aduk, Agl, Ak1, Ak2, Ak3, Ak6, Akt, AlaRS, Alat, alc, Ald1, Aldh, Alg13, Alk, alphaTub84B, AMPdeam, Amph, AMPKalpha, AnxB9, aPKC, apolpp, Apt1, Arc42, Archease, Argk1, Argk2, Arms, Arp1, Asator, Ask1, ASPP, AspRS, Atg1, Atg13, Atg17, Atpalpha, ATPsynB, ATPsynbeta, ATPsynCF6, ATPsynD, ATPsynF, ATPsynG, ATPsyngamma, ATPsynO, aurA, aurB, aux, awd, babo, ball, Bdbt, beta-PheRS, beta-Spec, betaTub56D, betaTub97EF, blw, bon, bsd, bsf, Bsg, bsk, bt, Btk, btl, Bub1, BubR1, by, cact, Cad96Ca, CaMKI, CaMKII, CanA-14F, CAP, Capr, capt, CASK, Cat, cbc, CCT2, CCT8, Cdc2rk, Cdc37, Cdc7, cdi, Cdk1, Cdk12, Cdk2, Cdk4, Cdk5, Cdk5alpha, Cdk7, Cdk8, Cdk9, Ced-12, Cerk, CG10177, CG10479, CG10513, CG10514, CG10550, CG10553, CG10559, CG10560, CG10562, CG10576, CG10602, CG10943, CG11089, CG11151, CG11350, CG11523, CG11594, CG11771, CG11811, CG11878, CG11889, CG11891, CG11892, CG11893, CG12016, CG12069, CG12147, CG1227, CG12289, CG12934, CG13360, CG13369, CG13474, CG13658, CG13659, CG13813, CG14207, CG14305, CG14314, CG14408, CG14505, CG15543, CG1648, CG1674, CG16898, CG16903, CG16985, CG17010, CG17237, CG17528, CG17597, CG17698, CG17896, CG1791, CG18765, CG2004, CG2201, CG2577, CG2794, CG3008, CG30178, CG30274, CG3107, CG31087, CG31097, CG31098, CG31099, CG31102, CG31104, CG31140, CG31145, CG31288, CG31300, CG31370, CG31380, CG31436, CG31624, CG31751, CG31832, CG31974, CG31975, CG31988, CG32195, CG3226, CG3277, CG32944, CG3301, CG33017, CG33098, CG33301, CG33509, CG33510, CG33511, CG34325, CG34384, CG3529, CG3534, CG3544, CG3603, CG3609, CG3630, CG3631, CG3699, CG3760, CG3902, CG40191, CG40635, CG41520, CG42258, CG42319, CG42366, CG4239, CG43078, CG43897, CG44085, CG44245, CG4461, CG4546, CG4629, CG4839, CG4945, CG5023, CG5126, CG5174, CG5446, CG5644, CG5757, CG5790, CG5828, CG6005, CG6650, CG6660, CG6800, CG6830, CG6834, CG6908, CG7094, CG7135, CG7156, CG7236, CG7328, CG7335, CG7409, CG7551, CG7920, CG8060, CG8173, CG8414, CG8441, CG8547, CG8565, CG8635, CG8841, CG9222, CG9259, CG9286, CG9391, CG9497, CG9498, CG9541, CG9593, CG9674, CG9784, CG9886, CG9961, CG9962, chic, chif, CHKov1, CHKov2, Chro, Cht9, cib, Cisd, CkIalpha, CkIIalpha, CkIIbeta, CkIIbeta2, ckn, Cks30A, Cks85A, Cmpk, colt, Coq8, cora, coro, corolla, Cortactin, COX4, COX5A, COX5B, cpb, cPges, Cpr, CRAT, Cs1, Csk, CycA, CycB, CycB3, CycC, CycD, CycE, CycG, CycH, CycJ, CycK, CycT, CycY, Cyp1, cype, Cyt-c1, Cyt-c-p, DAAM, dap, Dap160, dco, DCX-EMAP, Ddr, Dgk, Dgkepsilon, dia, Dic61B, dl, Dlat, Dld, dlg1, dnk, Doa, dokb, Dolk, dop, dos, Dpck, dpp, DppIII, Drak, Dref, Drep1, Droj2, Dsor1, Dyrk2, Dyrk3, eas, Eb1, Echs1, eEF1alpha2, eEF1gamma, eEF2, Egfr, Egfrap, eIF4B, eIF4E1, Eip63E, Elp3, Elp4, emb, Eno, Eph, Epp, eRF3, Erk7, ERp60, Etfb, Etf-QO, fab1, Fak, Fas1, fau, fax, fbl, FER, Fim, fj, Fkbp12, Fkbp59, flil, fln, fmt, for, fray, fs(1)h, fu, futsch, fwd, fwe, Gabat,

Galk, Galm1, gammaSnap1, Gapdh1, Gapdh2, GckIII, Gcn2, Gdh, gek, gish, Gk1, Gk2, Gldc, Gls, Glyp, GlyRS, Gnmt, gnu, Got2, Gpdh1, Gpo1, Gprk1, Gprk2, grk, grp, Grx3, Gs1, Gs2, gskt, GstD1, GstE12, GstO3, GstS1, GstT1, gwl, Had1, Haspin, Hecw, hep, Hex, Hex-A, Hex-C, Hexim, Hex-t1, Hex-t2, HHEX, Hil, Hipk, His1:CG33801, His4:CG33909, Hmbs, Hmgs, Hn, hop, hpo, hppy, Hrb27C, Hrs, Hsc70-1, Hsc70-2, Hsc70-4, Hsc70-5, Hsdl2, Hsp110, Hsp23, Hsp60B, Hsp83, htl, hts, I-2, Idgf3, Idgf6, Idh, Idh3a, Idh3b, Idh3g, if, IKKbeta, IKKepsilon, Ilk, inaC, inc, Indy, InR, IP3K1, IP3K2, Ip6k, Ipk1, Ipk2, Ire1, Irp-1A, Irp-1B, Ist1, Itgbn, jdp, jeb, JhI-26, JIL, JIL-1, kel, Klc, koko, KP78a, KP78b, ksr, kug, l(1)G0196, l(2)efl, l(2)gl, l(2)k01209, l(3)80Fj, LamC, LanB1, lap, Lar, Larp4B, Lasp, Ldh, lic, lig, LIMK1, Liprin-alpha, Lk6, Lkb1, Lmpt, lok, LqfR, Lrrk, Lsp1beta, LUBEL, mAcon1, mAcon2, Madm, Map205, MAPk-Ak2, Marc, Mat1, mats, mbc, mbt, Mcad, Mdh1, Mdh2, mEFTu1, mei, mei-41, Mekk1, meng, mew, Mf, Mhc, Mitofilin, Mkk4, Mlc1, Mlc2, Mlp60A, Mlp84B, mnb, Mo25, Mob2, Mob3, mod, Mos, Mp20, Mpcp1, Mps1, msn, Msp300, Mtch, mTor, Mtpalpha, Mulk, Mvk, mys, Myt1, NaCP60E, Nadk1a, Nadk1b, Nadk2, Nagk, Nak, NC2alpha, ND-20, ND-24, ND-39, ND-42, ND-49, ND-51, ND-75, ND-B14.5A, ND-B14.5B, ND-B16.6, ND-B17, ND-B17.2, ND-B22, ND-MLRQ, ND-SGDH, Nek2, Nepl21, niki, ninaC, nmd, nmdyn-D6, nmdyn-D7, nmo, Nmt, nonC, Nop56, Nplp2, Nrk, nrm, NTPase, Nuak, nudC, Obsc, Ogdh1, otk, p130CAS, p38a, p38b, p38c, p47, P5CDh1, P5cr-2, P5CS, Pak, Pak3, Papss, par, par-1, par-6, Parp16, parvin, Pask, Pax, pbl, Pcb, pcs, Pdha1, Pdhb, Pdk, Pdk1, Pdxk, PEK, Pepck2, Pfas, Pfdn6, Pfk, Pfrx, Pgam1, Pgam5, Pgi, Pgk, Pglym78, Pgm1, Phb2, PhKgamma, PhLP2, Pi3K21B, Pi3K59F, Pi3K68D, Pi3K92E, Pi4KIIalpha, Pi4KIIIalpha, Pink1, PIP4K, PIP5K59B, Pitslre, Pk34A, Pka, Pka-C1, Pka-C2, Pka-C3, Pka-R1, Pka-R2, Pkc53E, Pkc98E, Pkcdelta, PKD, Pkg21D, pkm, Pkn, Pli, pll, Pmvk, png, PNKP, polo, porin, Pp1-87B, Ppat-Dpck, Prm, Prosalpha1, Prosalpha1R, Prosalpha2, Prosalpha3, Prosalpha3T, Prosalpha4, Prosalpha4T1, Prosalpha4T2, Prosalpha5, Prosalpha6, Prosalpha6T, Prosalpha7, Prosbeta1, Prosbeta2, Prosbeta2R1, Prosbeta2R2, Prosbeta3, Prosbeta4, Prosbeta4R1, Prosbeta4R2, Prosbeta5, Prosbeta5R1, Prosbeta5R2, Prosbeta6, Prosbeta7, Prp4k, Prx3, Prx6c, Psi, Psn, Pstk, ptc, Ptth, Pur-alpha, put, Pvr, PVRAP, pyd3, Pyk, pzg, QIL1, Rab11, Rac1, Raf, Ran, Ras85D, rdgA, Rer1, Ret, RFeSP, Rfk, rhea, RIC-3, rictor, RIOK1, RIOK2, rl, Rok, Rop, Ror, RpL10Ab, RpL14, RpL23A, RpL40, RpLP1, Rpn1, Rpn10, Rpn11, Rpn12, Rpn12R, Rpn13, Rpn13R, Rpn2, Rpn3, Rpn5, Rpn6, Rpn7, Rpn8, Rpn9, RpS28b, RpS3A, Rpt1, Rpt2, Rpt3, Rpt3R, Rpt4, Rpt4R, Rpt5, Rpt6, Rpt6R, Rtnl1, S6k, S6kII, S6KL, SAK, salto, sax, scb, Scp1, Scsalpha1, ScsbetaA, scu, SdhA, SdhB, sdt, Sec6, Sema1a, sev, Sf3a1, sff, sgg, sgll, Shark, shep, shrb, Sik2, Sik3, Sin1, Sk1, Sk2, sktl, slgA, Slh, Slik, Slob, slpr, sls, smash, SmydA-2, SmydA-9, Snap29, SNF4Agamma, sni, Snrk, sns, Socs16D, Socs36E, Socs44A, spi, Spn43Ab, Sps2, sqa, Src42A, Src64B, SRPK, Srpk79D, Ssadh, Ssl, sta, Stam, stck, Ste12DOR, Ste:CG33237, Ste:CG33245, Ste:CG33239, Ste:CG33236, Ste:CG33240, Ste:CG33244, Ste:CG33241, Ste:CG33247, Ste:CG33246, Ste:CG33243, Ste:CG33238, Ste:CG33242, SteXh:CG42398, sti, Stim, Stip1, Stlk, Strn-Mlck, STUB1, stv, su(r), Tab2, Taf1, Tak1, Takl1, Takl2, Tango10, Tao, Tcs5, Tctp, tefu, Ten-m, TER94, ThrRS, Tie, Tkt, tkv, Tlk, Tm1, Tm2, tmn, tmod, tn, tor, Tpi, TpnC4, Tps1, trbl, trc, trk, trsn, Trx2, Tsp42Ea, tsr, Tssk, Ttc1, Txl, tzn, Uba1, Uch-L5, Uch-L5R, Uck, Ugt49B1, Unc-115a, unc-45, uncNacbeta, up, UQCR-14, UQCR-C1, UQCR-C2, Vajk4, ValRS, vari, Vha100-1, Vha100-2, Vha13, Vha26, Vha55, Vha68-1, Vha68-2, VhaM9.7-b, vig, Vinc, Vps15, Vps60, wdb, Wee1, wit, wnd, Wnk, wrd, Wsck, wts, wupA, yki, Yp1, Yp3, Zasp52, Zasp66, Zasp67, zip, zormin, Zyx

**Supplementary text 2. List of Ig19 ligands with models CPAE below 10.**

Abl, Ack, Ack-like, Adk1, Adk3, Aduk, Ak1, alc, Alk, apolpp, Arms, Asator, ASPP, Atg1, Atg13, Atg17, ATPsynG, bon, bsd, bt, btl, BubR1, cact, Cad96Ca, Capr, capt, CASK, cbc, Cdc37, cdi, Cdk12, Cdk8, Cdk9, Cerk, CG10177, CG10514, CG10553, CG10560, CG10562, CG11350, CG11878, CG11892, CG11893, CG12147, CG12934, CG13360, CG13659, CG14314, CG14408, CG15543, CG1674, CG16898, CG16903, CG17698, CG30274, CG31087, CG31098, CG31099, CG31102, CG31104, CG31140, CG31300, CG31436, CG31832, CG31974, CG32195, CG33017, CG33301, CG34384, CG3603, CG3760, CG3902, CG40191, CG41520, CG43078, CG43897, CG44085, CG44245, CG4546, CG4839, CG4945, CG5790, CG6005, CG6660, CG7094, CG8441, CG8547, CG8635, CG9259, CG9497, CG9498, CG9541, CG9674, CG9961, chif, CHKov2, CkIIbeta, CkIIbeta2, ckn, cpb, CRAT, Csk, CycA, CycB3, CycC, CycE, CycT, CycY, Cyp1, Cyt-c1, DAAM, Dap160, dnk, dop, dos, dpp, Drep1, Dtymk, Dyrk2, Dyrk3, Egfrap, eIF4B, Eip63E, Elp4, Eno, Erk7, fab1, Fak, fbl, FER, fliI, fmt, for, fray, fs(1)h, futsch, fwd, GckIII, gek, gish, gnu, Gprk2, grp, gskt, GstE12, gwl, Hecw, hep, Hex, Hexim, Hil, Hipk, His1, hop, hppy, Hrb27C, Hrs, Hsc70-2, Hsc70-5, Hsc70Cb, Hsdl2, Hsp110, Hsp60B, htl, hts, Hykk, Idgf6, if, inc, Indy, InR, IP3K2, Ip6k, Ipk1, jdp, JIL, JIL-1, kel, KP78b, ksr, kug, l(1)6Fd, l(2)gl, l(3)80Fj, LamC, Lar, Larp4B, Lasp, LIMK1, Liprin-alpha, Lk6, Lmpt, LqfR, Lrrk, LUBEL, Madm, Map205, mats, mbc, Mcad, mei, Mekk1, mew, Mhc, Mkk4, Mlc2, mnb, Mob2, Mps1, msn, Msp300, Mulk, mys, Myt1, NaCP60E, Nadk1a, Nadk1b, Nadk2, Nak, NC2alpha, ND-39, ND-49, ND-B14.5A, nmd, nmo, Nol9, nonC, Nuak, Obsc, Ogdh1, oya, p130CAS, p47, Pak, Papss, Pask, Pax, pbl , Pdk, Pdk1, PEK, Pfk, Pfrx, Pgk, PhKgamma, Pi3K21B, Pi3K68D, Pi3K92E, Pi4KIIalpha, PIP5K59B, Pka, PKD, pkm, Pli, porin, Prm, Prosalpha1, Prosalpha1R, Prosalpha2, Prosalpha3, Prosalpha3T, Prosalpha4, Prosalpha4T1, Prosalpha5, Prosalpha6T, Prosbeta1, Prosbeta2, Prosbeta2R2, Prosbeta3, Prosbeta4R2, Prosbeta6, Prosbeta7, Prp4k, Psn, ptc, Pvr, Rab11, Ran, rdgA, Ret, rhea, RIC-3, rictor, RIOK2, Rpn12, Rpn2, Rpn6, Rpt1, Rpt2, Rpt3, Rpt3R, Rpt5, Rpt6, Rpt6R, Rtnl1, S6k, S6kII, scb, sdt, Sema1a, sev, sff, sgg, sgll, shep, shrb, Sik2, Sik3, sktl, Slik, Slob, slpr, sls, smash, SNF4Agamma, Snrk, sns, Socs16D, Socs36E, spi, Src42A, SRPK, Srpk79D, Ssl, Stam, Strn-mck, STUB1, stv, Tab2, Taf1, Tak1, Tao, Ten-m, TER94, Tie, Tlk, tn, Tps1, trbl, trc, trsn, Tsp42Ea, Uba1, Uck, Unc-115a, Vha100-2, Vha26, Vha68-1, Vha68-2, vig, Vps60, wall, Wee1, wit, wnd, Wsck, wts, wupA, yki, Yp3, Zasp52, Zasp66, Zasp67, zip, zormin

**Supplementary text 3. List of Ig19 ligands with models CPAE below 5.**

Ack-like, apolpp, Atg17, bt, cact, capt, cdi, Cdk12, CG12934, CG14408, CG16898, CG31098, CG31436, CG31974, CG33017, CG33301, CG43897, CG44245, CG6660, CG9259, chif, CycE, CycY, dop, dos, dpp, Dyrk3, Egfrap, Eip63E, fab1, futsch, fwd, GckIII, hep, Hexim, Hil, Hsc70Cb, Hsp110, hts, KP78b, l(3)80Fj, Larp4B, LIMK1, Lk6, LqfR, Mcad, mnb, Mob2, Msp300, Mulk, ND-49, Nuak, p130CAS, Pak, Pax, PhKgamma, Pi3K68D, Prm, Prosalpha4T1, Prosbeta3, Prosbeta4R2, Psn, ptc, rhea, RIC-3, Rpt3R, Rtnl1, sdt, sff, Sik3, Slik, Slob, sls, SNF4Agamma, Socs16D, Src42A, Srpk79D, Tab2, Tlk, tn, Tps1, Uba1, Vha68-2, Zasp66

**Supplementary text 4. Amino acid sequences of Cher Ig domains used.**
Cher Ig18-Ig19:
AGEGSNRKREKIQRERDAVPITEIGSQCKLTFKMPGITSFDLAACVTSPSNVTEDAEIQEVEDGLYAVHFVPKE
LGVHTVSVRYSEMHIPGSPFQFTVGPLRDSGSHLVKAGGSGLERGVVGEAAEFNVWTREAGGGSLAISVEG
PSKADIEFKDRKDGSCDVSYKVTEPGEYRVGLKFNDRHIPDSPFKVYV
Cher Ig16-Ig17:
TGEGRKRNQQSVGSCSEVTMPGDITDDDLRALNASIQAPSGLEEPCFLKRMPTGNIGISFTPREIGEHLVSV
KRLGKHINNSPFKVTVCEREVGDAKKVKVSGTGLKEGQTHADNIFSVDTRNAGFGGLSVSIEGPSKAEIQCT
DKDDGTLNISYKPTEPGYYIVNLKFADHHVEGSPFTVKV
Cher Ig19:
LRDSGSHLVKAGGSGLERGVVGEAAEFNVWTREAGGGSLAISVEGPSKADIEFKDRKDGSCDVSYKVTEPG
EYRVGLKFNDRHIPDSPFKVYV
Cher Ig17:
CEREVGDAKKVKVSGTGLKEGQTHADNIFSVDTRNAGFGGLSVSIEGPSKAEIQCTDKDDGTLNISYKPTEP
GYYIVNLKFADHHVEGSPFTVKV

**Supplementary text 5. R code to prepare aminoacid sequences for Colabfold.**

```
# Load required library
library(seqinr)
# Read a FASTA file containing all potential CHER-interacting proteins
f <- read.fasta("X.fasta", seqtype = "AA")
# Define the Ig19 sequence
Ig19 <-
"LRDSGSHLVKAGGSGLERGVVGEAAEFNVWTREAGGGSLAISVEGPSKADIEFKDRKDG
SCDVSYKVTEPGEYRVGLKFNDRHIPDSPFKVYV"

# Function to splits sequences into 200-amino-acid fragments  and append Ig19
ff <- vector()  # Initialize a vector to store processed sequences
for (i in 1:length(f)) {
 q <- unlist(getSequence(f[i], as.string = FALSE))  # Extract sequence
 n <- getName(f[i])  # Get sequence name
 w <- split(q, ceiling(seq_along(q) / 200))  # Split sequence into chunks of 200 amino
acids
 for (ee in 1:length(w)) {
  # Concatenate sequence chunks and append Ig19
  ff <- c(ff, paste(paste(unlist(w[ee]), collapse = ""), ":", "Ig19", sep = ""))
 }

 # Write sequences to individual FASTA files
 write.fasta(f[i], n, paste0(n, ".fasta"))
```

```
 # Write processed sequences with Ig19 to FASTA files
 for (ee in 1:length(ff)) {
  write.fasta(ff[ee], n, paste0(n, "_Ig19.fasta"))
 }
}
```

**Supplementary text 6. R code to find the contact sites and calculate the CPAE values from ColabFold models.**

```
#install Bio3D and jsonlite packages into R
install.packages("bio3d")
install.packages("jsonlite")

# Load the libraries
library(bio3d)
library(jsonlite)

# Function that finds the binding sites and gives the PAE of the binding site
predict_alignment_error <- function(pdb_file, json_file) {
 # Read the PDB file and the JSON file
 pdb <- read.pdb(pdb_file)
 json_data <- fromJSON(json_file)
 # Extract the two chains from the PDB object
 chain1 <- trim.pdb(pdb, chain="A")
 chain2 <- trim.pdb(pdb, chain="B")
 # Identify the binding site in chain1 and retrieve the residue numbers
 bs1 <- binding.site(chain1, chain2, cutoff=5)
 # Gives a vextor of interacting residues numbers
 bs1_resno <- bs1$resno
 # Identify the binding site in chain2 and retrieve the residue numbers
 bs2 <- binding.site(chain2, chain1, cutoff=5)
 # Gives a vextor of interacting residues numbers
 bs2_resno <- bs2$resno
 # Extract the predicted alignment error between Chain A and Chain B
 chain1_l <- max(chain1$atom$resno)
 mat <- json_data$pae
 mean_error <- mean(c(mat[bs1_resno,
c(chain1_l+bs2_resno)],mat[c(chain1_l+bs2_resno),bs1_resno]))
 return(list(mean_error=mean_error, bs1_resno=bs1_resno,
bs2_resno=bs2_resno))
}

# List all files in the folder
```

```
all_files <- list.files()

# Filter files that have either .pdb or .json extension
pdbs <- all_files[grep("\\.pdb$", all_files)]
jsons <- all_files[grep("\\000.json$", all_files)]

# run in a loop for all the files in the folder
pae=vector()
for (i in 1:length(pdbs))
  {pae[i]=predict_alignment_error(pdbs[i],jsons[i])}
```

### Supplementary Figures

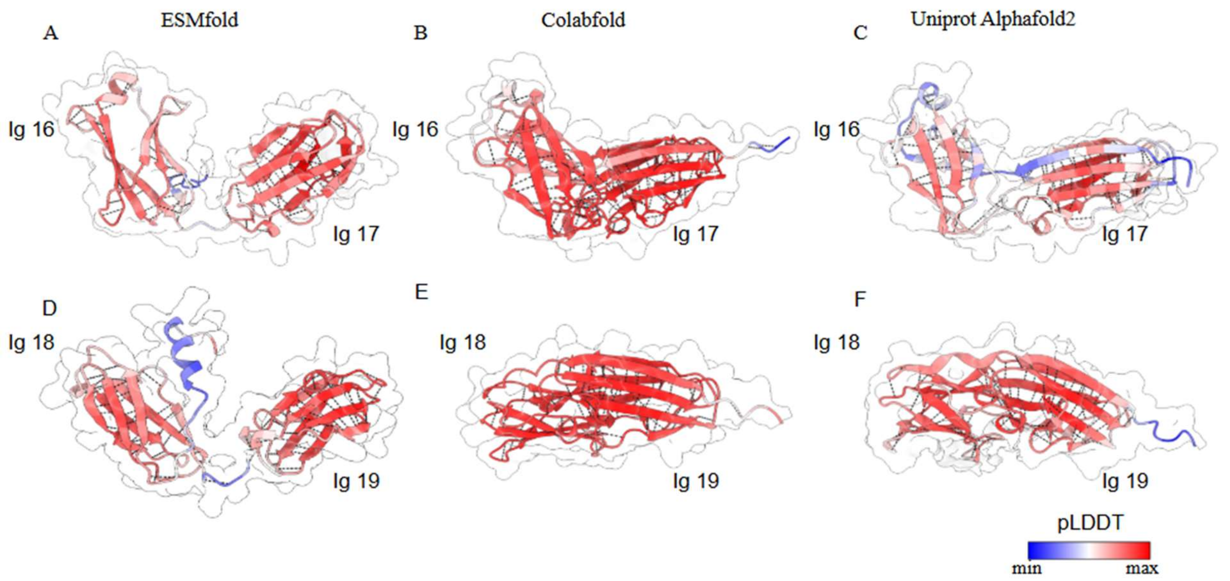


**Supplementary Figure 1. pLLDDT values in Ig16-Ig17 and Ig18-Ig19 models. A)** Predicted model of Ig16 and Ig17 using ESMFold. **B)** Predicted model of Ig16 and Ig17 using ColabFold. **C)** Predicted model of Ig16 and Ig17 using AlphaFold 2, obtained from UniProt. **D)** Predicted model of Ig18 and Ig19 using ESMFold. E**)** Predicted model of Ig18 and Ig19 using ColabFold. **F)** Predicted model of Ig18 and Ig19 using AlphaFold 2, obtained from UniProt.

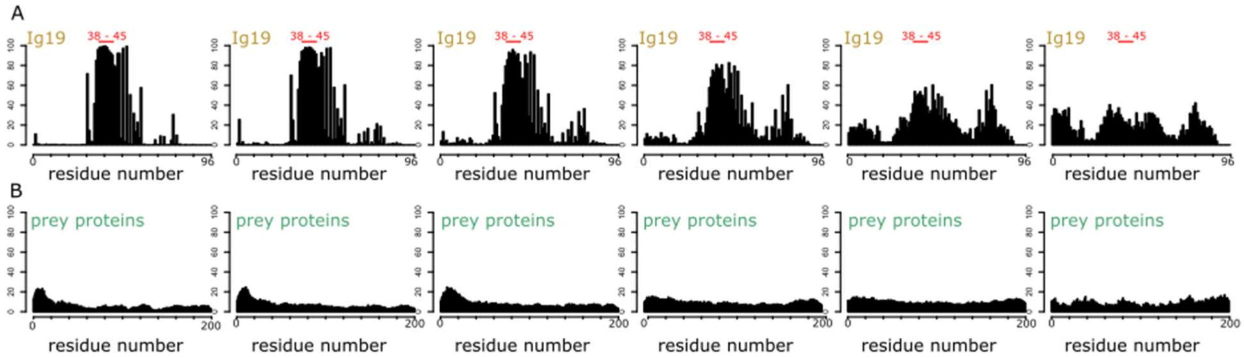


**Supplementary Figure 2. Histograms of contact sites between Ig19 and the prey proteins, separated by different CPAE scores.** (A) Histograms of the contacts in Ig19 show a clear enrichment of binding events toward the expected strand, located at positions 38–45. (B) Histograms of the binding events in the prey proteins reveal a slight preference at the N-terminal end, though no clear enrichment is observed otherwise. In both A and B, the CPAE scores for each plot are, from left to right: 0–5, 5–10, 10–15, 15–20, 20–25, and 25–30.

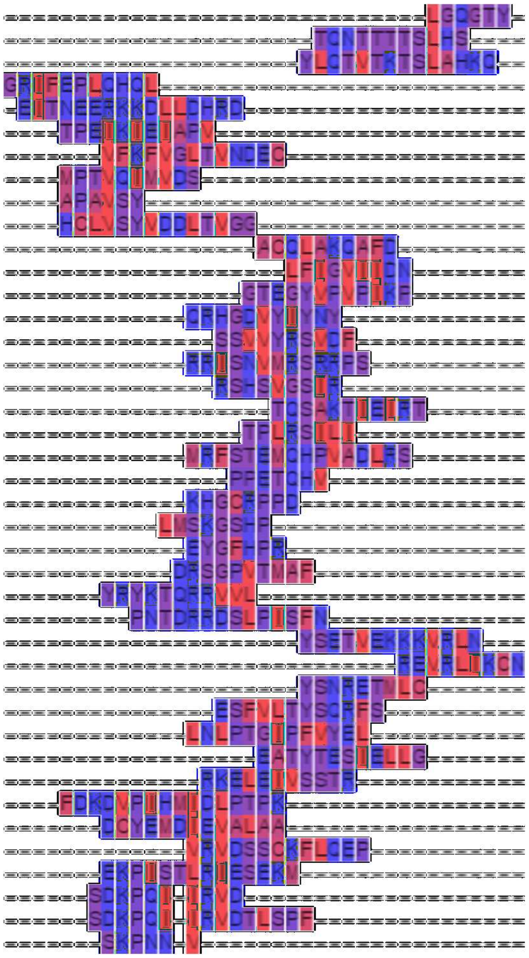

**Supplementary Figure 3. Multiple sequence alignment from the contact sites of all models with CPAE values below 5 to Ig19.**
